# Number of kinesins engaged in axonal cargo transport: A novel biomarker for neurological disorders

**DOI:** 10.1101/2023.08.03.551910

**Authors:** Kumiko Hayashi, Kazuo Sasaki

**Affiliations:** Institute for Solid State Physics, The University of Tokyo, Kashiwa, Japan; Department of Applied Physics, Graduate School of Engineering, Tohoku University, Sendai, Japan

**Author notes:** Correspondence to Kumiko Hayashi.

## Abstract

Kinesin motor proteins play crucial roles in anterograde transport of cargo vesicles in neurons, moving them along axons from the cell body towards the synaptic region. Not only the transport force and velocity of single motor protein, but also the number of kinesin molecules involved in transporting a specific cargo, is pivotal for synapse formation. This collective transport by multiple kinesins ensures stable and efficient cargo transport in neurons. Abnormal increases or decreases in the number of engaged kinesin molecules per cargo could potentially act as biomarkers for neurodegenerative diseases such as Alzheimer’s, Parkinson’s, amyotrophic lateral sclerosis (ALS), spastic paraplegia, polydactyly syndrome, and virus transport disorders. We review here a model constructed using physical measurements to quantify the number of kinesin molecules associated with their cargo, which could shed light on the molecular mechanisms of neurodegenerative diseases related to axonal transport.

## Introduction

In neurons, anterograde transport is characterized by the trafficking of synaptic materials, synthetized in the cell body, along axons and up to the synaptic terminals by kinesin motor proteins. Kinesin is a motor protein that processively moves mainly in the plus-end direction of microtubules by obtaining energy through ATP hydrolysis (Hirokawa et al., 2009; Vale, 2003). In contrast, the molecular motor known as dynein moves towards the minus end of the microtubule (Vale, 2003). Dynein functions to transport cargo in the direction opposite to that of kinesin, heading towards the microtubule’s minus end. This allows for efficient multi-directional transport within the cell. Dynein and kinesin work in concert to facilitate complex intracellular transport processes. It is known that there are 45 types of kinesin superfamily members in mammalians such as humans and mice (Hirokawa et al., 2010). Among them, KIF1A is a kinesin that transports synaptic vesicle precursors, and it is known that mutations in KIF1A can cause excessive transport of synaptic vesicle precursors and transport disorders, leading to abnormalities in synapse formation (Anazawa et al., 2022; Chiba et al., 2019; Niwa et al., 2016). As a result, the mutations in KIF1A lead to severe neurodegenerative diseases (Gabrych et al., 2019; Guedes-Dias and Holzbaur, 2019). While the genetic diseases caused by KIF1A are rare (Iqbal et al., 2017; Klebe et al., 2012), a significant number of people are suffering from them, and a global community to support those with KIF1A-related neurodevelopmental disorders, KIF1A.org, was established in 2016.

Axonal transport deficits cause by KIF1A mutations have been linked to physical deficits, such as a decrease in transport velocity, transport force, or the attachment rate to microtubules (Anazawa et al., 2022; Budaitis et al., 2021; Chiba et al., 2019; Lam et al., 2021). In addition, we have discovered that changes in the number of kinesin molecules transporting cargo can disrupt synapse formation (Hayashi et al., 2018a). It is thought that axonal transport is efficient if the cargo (synaptic vesicle precursors) is transported by the appropriate number of KIF1A molecules, and becomes inefficient when the number of KIF1A molecules decreases or increases excessively (Hayashi et al., 2018a).

We will discuss a novel measurement method to estimate the number of kinesins, engaging in the transport of synaptic vesicle precursors in the motor neurons of *Caenorhabditis elegans* (*C. elegans*) as an example. Figure 1 shows the results of observing the transport of GFP-labeled synaptic vesicle precursors in the axon using fluorescence microscopy. We can observe the cargo vesicles transported by kinesins, moving in one direction while fluctuating (tracked by particle tracking software). The causes of this fluctuating behavior are thermal fluctuations and collisions with surrounding vesicles and the cytoskeleton.

**Fig. 1.**
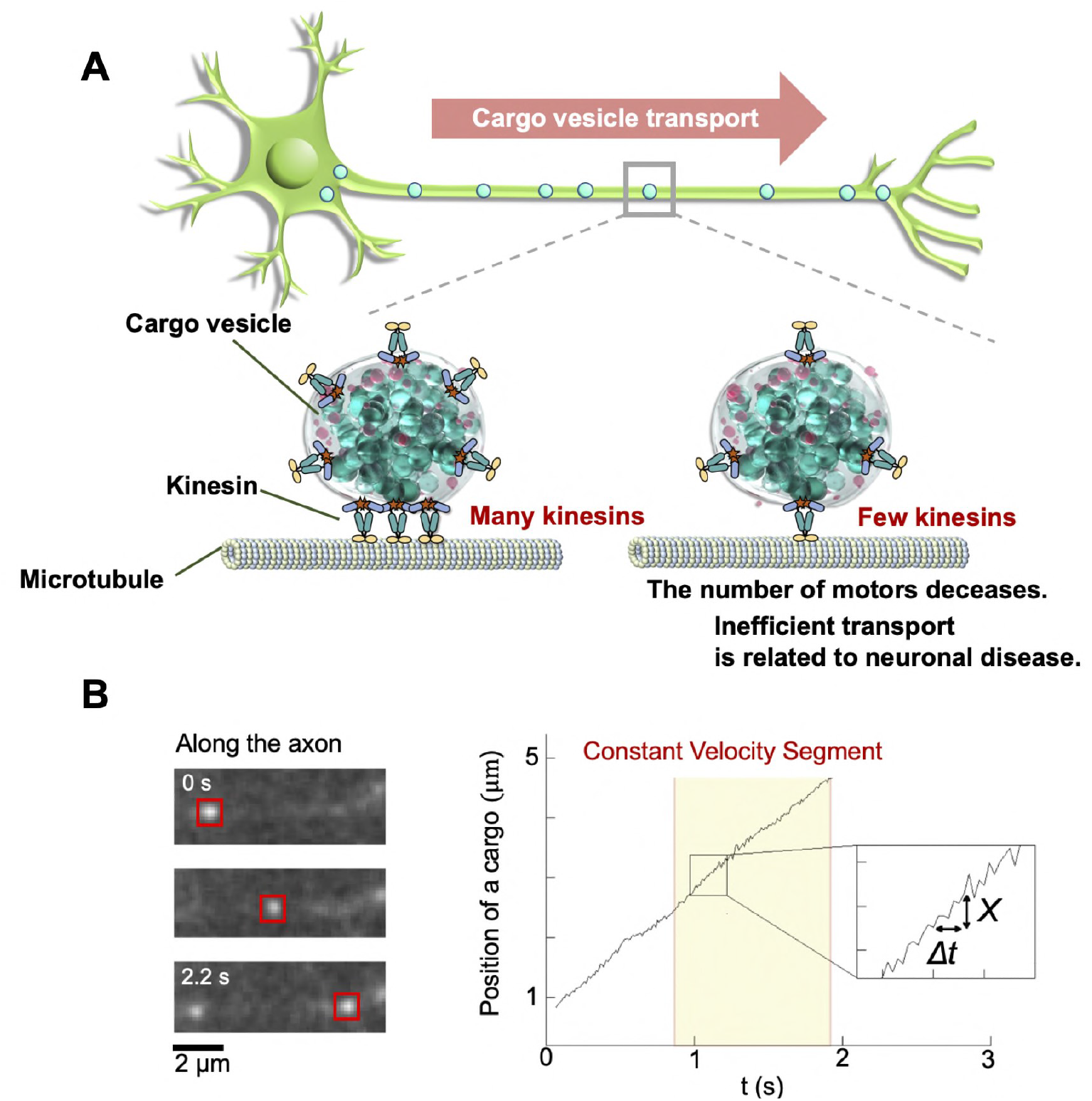
A: Schematics of cargo vesicle transport by kinesin. If the number of kinesins is appropriate, neuronal axonal transport remains stable. However, if the number decreases, it can lead to neurodegenerative diseases. **B:** Fluorescence observation of a cargo vesicle, as reported by (Hayashi et al., 2018a)(left). The image tracks the position of the cargo over time. The inset represents *X* and Δ*t*, while the highlighted yellow segment represents a constant velocity segment.

The details of the physical background of the measurement method are left to the biophysical references (Hasegawa et al., 2019; Hayashi et al., 2018a; Hayashi et al., 2021; Hayashi et al., 2018b), we have successfully identified the parameter χ, a potential biomarker indicative of the quantity of kinesin molecules associated to a single cargo. Measuring the parameter χ, we found that 1-6 kinesin molecules are required to transport synaptic vesicle precursors. Higher number of kinesin molecules leads to a more efficient transport of the cargo while lower number of kinesins and weak binding are associated with a deficit in axonal transport (Hayashi et al., 2018a).

### Theoretical model of cargo transport to derive χ

Let’s denote the displacement of a cargo vesicle during the time interval Δ*t* as *X*, and the number of kinesins transporting a single cargo vesicle as *N* (Fig. 2A). *x*_*i*_ is the displacement of the *i*-th kinesin (*i* = 1,2, ⋯, *N*). Then, the probability distribution function of the displacement *X* of the cargo vesicle can be written as:

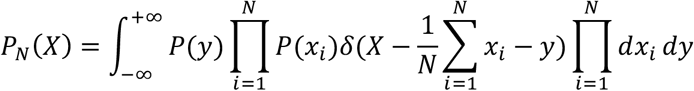

By the definition of the delta function

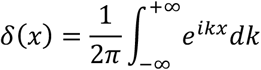

we have:

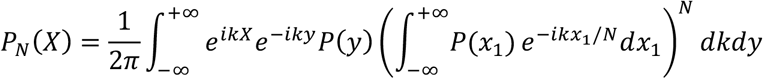

Here, we assume that the cargo vesicle surface (membrane) is flexible and each kinesin moves independently (Fig. 2A). *P*(*x* _*i*_) is written as

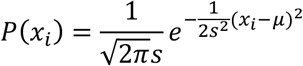

Here 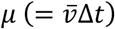 and *s*^2^ (= *σ*^2^Δ*t*) are the displacement and variance of each kinesin, respectively. *P*(*y*) represents the effect of thermal noise acting on the cargo vesicle (Fig. 2A),

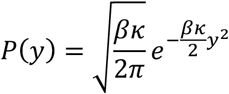

where *κ* is an effective stiffness between the cargo and kinesin molecules, and *β* = *k*_B_*T* (*k*_B_ :Boltzmann constant, *T*:Temperature of the environment (temperature in a cell)). Substituting the two equations into *P*_*N*_(*X*), we obtain

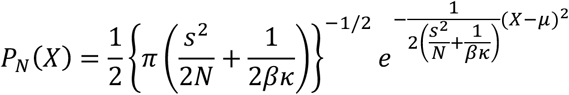

When considering a sufficiently large Δ*t* (Δ*t* → ∞) and since *s*^2^ ∝ Δ*t, s*^2^/*N* ≫ 1/*βκ* (= const), we have:

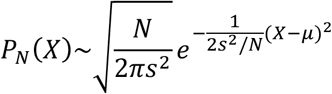

At this point, we focus on the mean 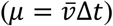 and variance (2*D*Δ*t* = *s*^2^/*N*) of the cargo vesicle displacement *X*, which are measurable quantities 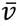 and D in the experiment, and construct the following quantity:

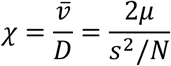

In Fig. 2B, we provide the simplest force (load)-velocity (*F*-*v*) relationship model of kinesin. It should be noted that there exist more sophisticated models of the force-velocity relationship, such as the energy landscape model (Schnitzer et al., 2000) and the three-state model (Sasaki et al., 2018), both of which are derived based on the ATP hydrolysis mechanism of kinesin. The *F*-*v* relationship was measured in the *in-vitro* single-molecule experiments (Nishiyama et al., 2002; Schnitzer et al., 2000).

**Fig. 2.**
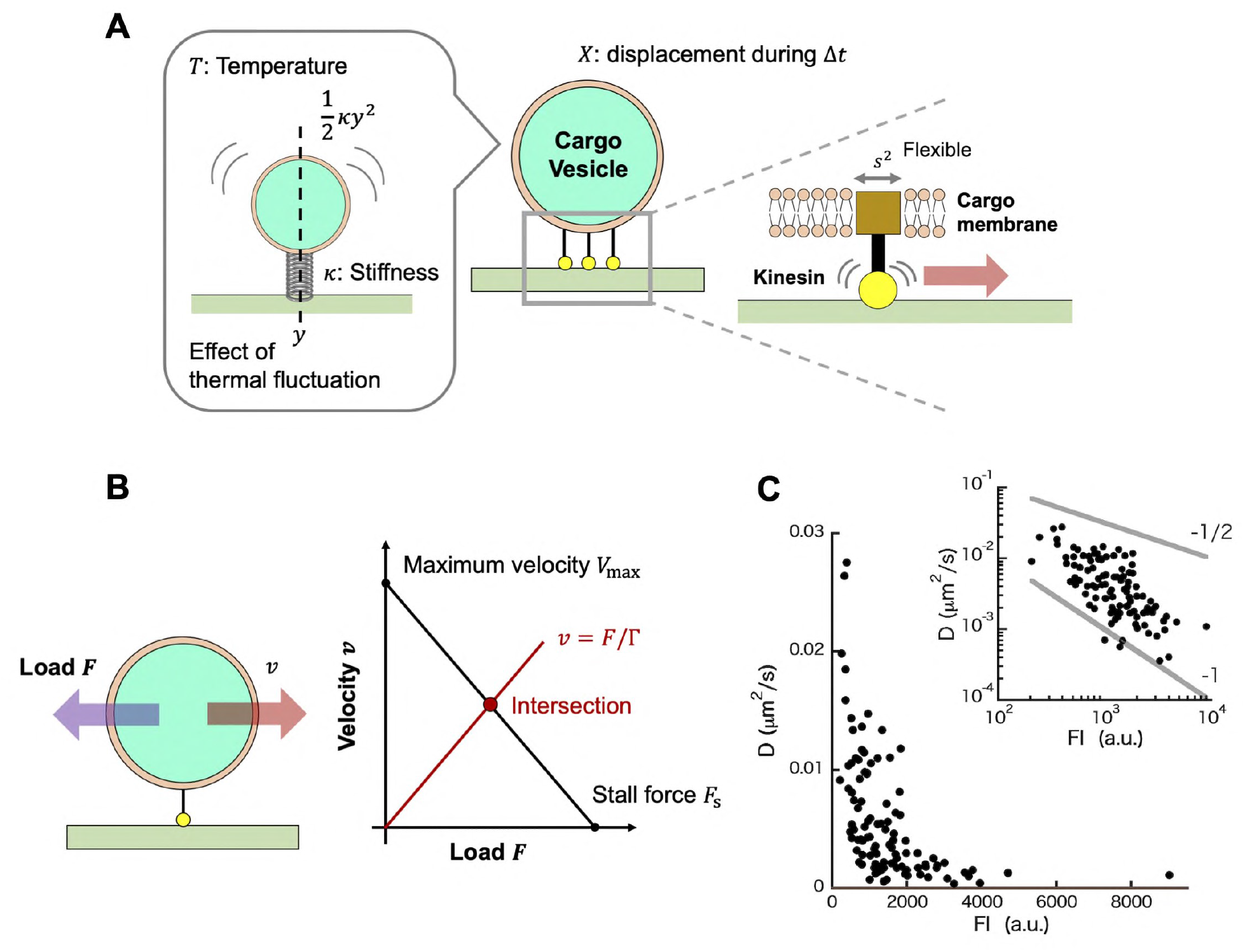
A: Theoretical model of cargo vesicle transport by kinesin. **B:** Simple force-velocity relationship of kinesin. **C:** *D*(= σ^2^/2*N*Δ*t*) versus the fluorescence intensity (FI) of cargo vesicles observed in the mouse hippocampal neurons, originally obtained in the supporting material of the reference (Hayashi et al., 2018a).

When the viscosity coefficient of the vesicle is denoted by Γ, the average velocity of the cargo vesicle 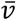 can be written as the intersection between the force-velocity (*F*-*v*) relationship and the line representing the Stokes law (*v* = *F*/Γ) as follows:

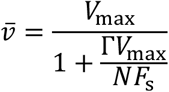

where *V*_max_ and *F*_s_ is the maximum velocity (the velocity without a cargo vesicle) and the maximum force (the stall force that kinesin can generate against a load *F*). *F*_s_ is known to be of the order of 10 pN from the single-molecule experiments using optical tweezers (Nishiyama et al., 2002; Schnitzer et al., 2000), and *V*_max_ is reported to be 4 *μ*m/s as a result of the extreme-value analysis (Naoi et al., 2021). From the result of fluid mechanics, Γ = 6π*ηr* (*η* :viscosity of the environment, *r* :radius of the cargo). Then, the expression of χ including the vesicle’s viscosity coefficient Γ is:

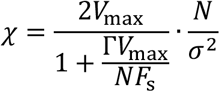

Note that 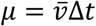 and *s*^2^ = *σ*^2^Δ*t*. The *N* dependence of χ is explicitly written as

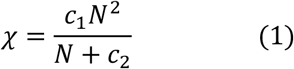

where *c*_1_ = 2*V*_max_/*σ*^2^ and *c*_2_ =Γ*V*_max_/*F*_s_. Thus, the number of kinesin molecules *N* can be related to χ, a measurable quantity in the experiment.

Under the high viscosity inside the cell 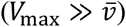, where we have:

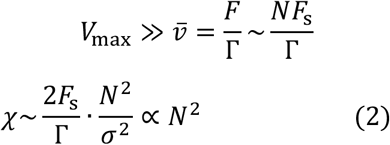

As to the variance *σ*^2^ of a kinesin, the fluorescence observation of cargos in neurons (Hayashi et al., 2021) in Fig. 2C suggests 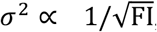, where FI is the fluorescence intensity of a cargo vesicle, i.e., 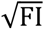 represents the cargo radius when the fluorescence materials are distributed on the cargo vesicle surface uniformly. It indicates *σ*^2^ ∝ 1/Γ because Γ = 6π*ηr* . Combine the experimental suggestion with Eq. (2), the following relation is predicted,

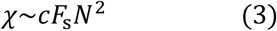

Here *c* is the constant which has a unit of energy such as *k*_B_*T* . It is a future research to derive Eq. (3) theoretically.

### Experimental results

We show the experimental results to measure χ (Eq. (1)). Fluorescence images of green fluorescence protein (GFP)-labeled synaptic cargo (synaptic vesicle precursors) transported by kinesin, UNC104/KIF1A, in the axon of *C. elegans* were captured using a camera at 100 frames per second as described in Fig. 1B (refer to the reference (Hayashi et al., 2018a) for the detailed experimental procedures). The body movements of *C. elegans* worms were suppressed under anesthesia (Niwa, 2017).

We consider a theoretical model for the motion of a cargo vesicle, transported by kinesins in constant velocity segments (CVSs) (Fig. 1B), which is assumed to be unaffected by other motor proteins based on previous experimental studies (D’Souza et al., 2022; Gennerich and Schild, 2006). This implies that the tugofwar model, depicting a conflict between the two motors moving in opposite directions, kinesin and dynein (Gennerich and Schild, 2006; Gross, 2004), was not considered for CVSs. Instead, we support the ‘motor coordination model,’ in which adaptor proteins that connect motors with cargos deactivate opposing motors (Chen et al., 2019).

For one such segment, we computed χ as a function of Δ*t*, and observed that χ converges to a constant value as Δ*t* becomes large (Fig. 3A). Subsequently, we calculated the χ values for 40 different cargo vesicles and observed the discrete behavior of χ (Fig. 3B, left). We clustered χ based on its discreteness, and then plotted χ against the cluster number *N*. We subsequently examined the dependency of χ on *N*, as predicted by Eq. (1) (Fig. 3C).

**Fig. 3.**
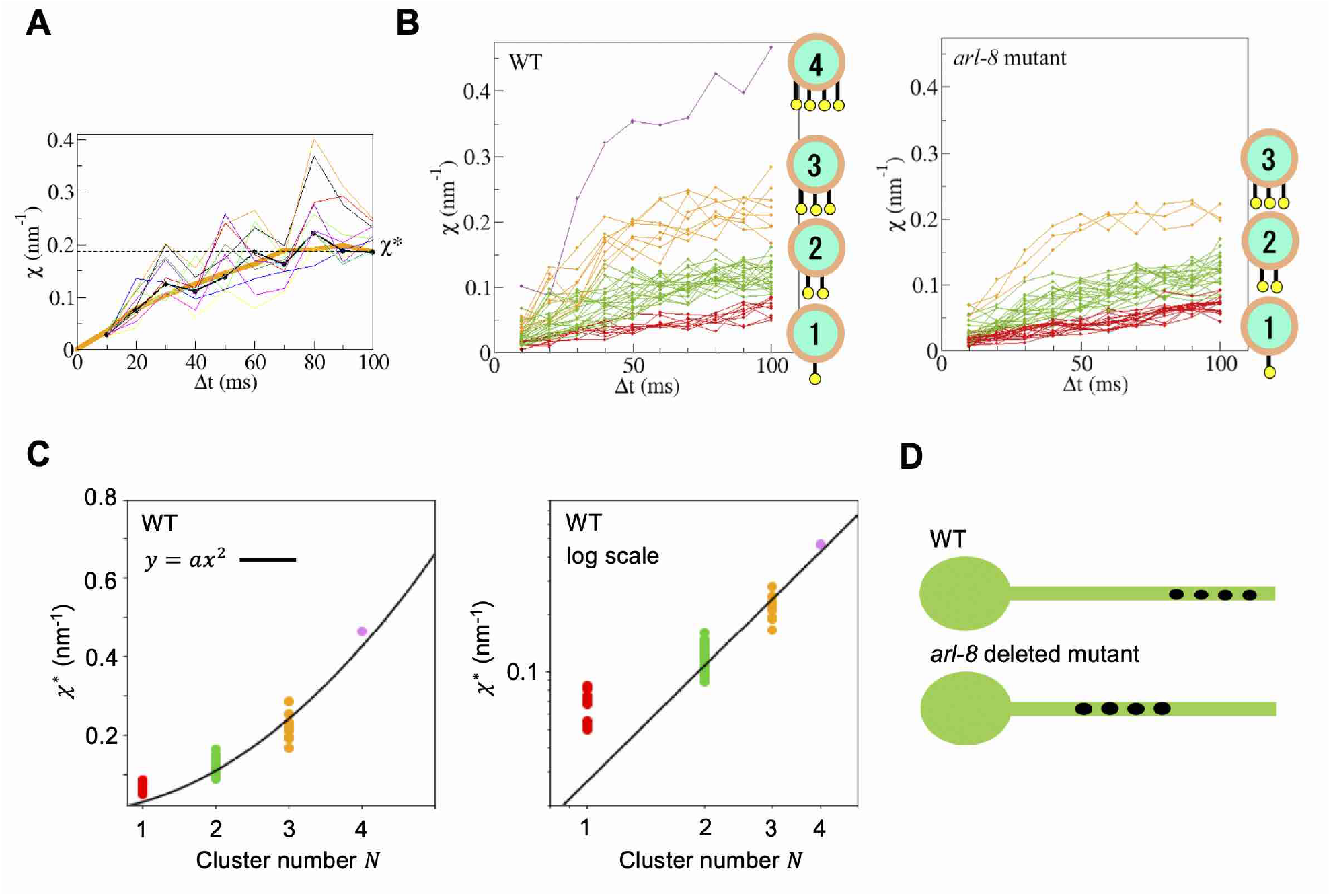
Measurement of χ for synaptic vesicle precursor transport by UNC104/KIF1A in *C. elegans* motor neurons (Hayashi et al., 2018a). **A:** Measurement of χ for a cargo vesicle as an example. The thin curves represent χ calculated based on a boot strapping method for the error estimation (Hayashi et al., 2018a). **B:** Measurement of χ for 40 cargo vesicles in the case of the wild type *C. elegans* (left) and the *arl-8* deleted mutant *C. elegans* (right). We found four clusters for the wild type and three clusters for the mutant. **C:** The convergent value, χ^∗^, of χ as a function of the cluster number *N*. The black line represents Eq. (2). **D:** Schematic of *en passant* synapses in the motor neuron of *C. elegans*.

Kinesins that are not engaged in vesicle transport exist in an autoinhibitory (inactive) state to prevent ATP wastage. Previous studies revealed that the protein ARL-8 promotes the release of kinesin from this autoinhibitory state (Niwa et al., 2016). In *arl-8* deleted *C. elegans* mutants, the release of kinesin autoinhibition does not proceed adequately, leading to a decrease in transport stability due to a shortage of active kinesin molecules. This causes abnormalities in synapse formation such as mis-locations of synapses (Fig. 3D) (Niwa et al., 2016). We performed the measurement of χ using the same *arl-8* deleted *C. elegans* mutant (Hayashi et al., 2018a). The results are shown in Fig. 3B (right). Consequently, we found a decrease in the number of χ clusters, indicating that χ could detect the reduction in the number of active kinesin molecules.

We also measured χ in mouse neurons (Hayashi et al., 2021; Hayashi et al., 2018b). In mouse hippocampal neurons, 1-6 kinesin molecules were identified to be involved in the transport of synaptic cargos (Fig. 4A and B). In mouse superior cervical ganglion (SCG) neurons, we found 1-4 kinesin molecules to be involved in the transport of endosomes.

**Fig. 4.**
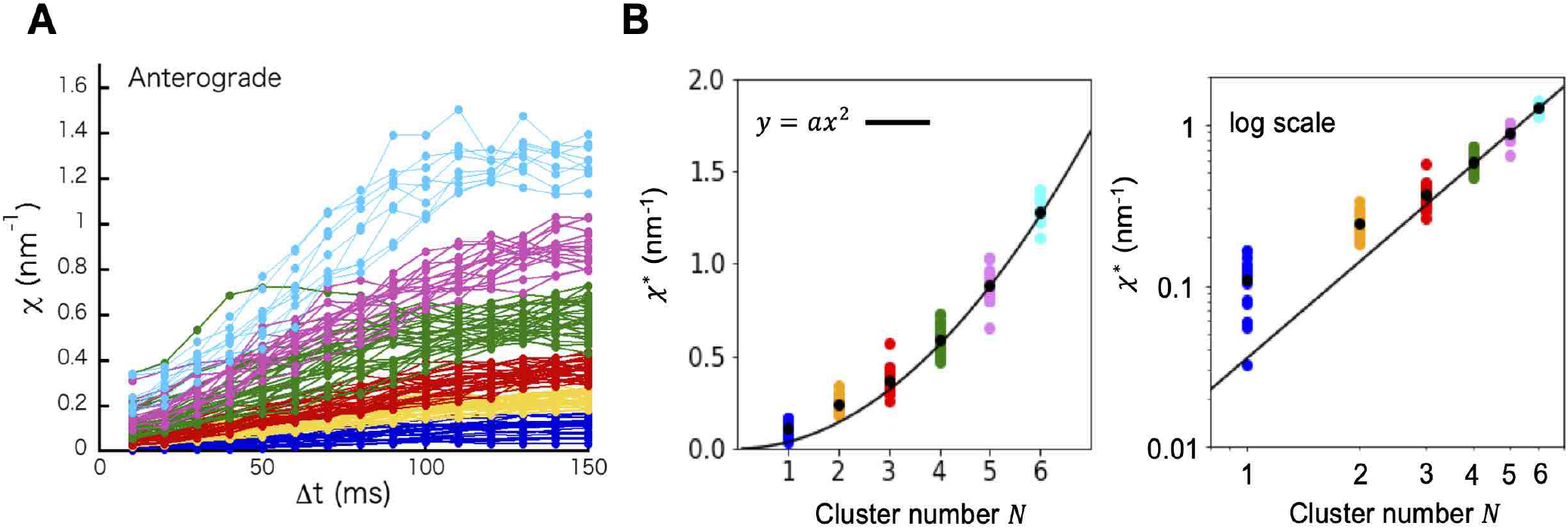
Measurement of χ for synaptic vesicle precursor transport by KIF1A in mouse hippocampal neurons (Hayashi et al., 2021). **A:** Measurement of χ for 131 cargo vesicles. We found six clusters. **B:** The convergent value, χ^∗^, of χ as a function of the cluster number *N*. The black line represents Eq. (2).

Regarding dynein, another motor protein, which moves in opposite direction of kinesin (Guedes-Dias and Holzbaur, 2019), previous studies have reported on the measurement results of χ (Hasegawa et al., 2019; Hayashi et al., 2021; Peng et al., 2022) (Fig. 5). The different structures and *F*-*v* relationships have been reported for dynein (Brenner et al., 2020; Elshenawy et al., 2019; Gennerich et al., 2007; Hirakawa et al., 2000; Mallik et al., 2004; Toba et al., 2006). Retrograde transport by dynein is also known to be related to neurodegenerative diseases (Guedes-Dias and Holzbaur, 2019; Schmieg et al., 2014; Terenzio et al., 2017).

**Fig. 5.**
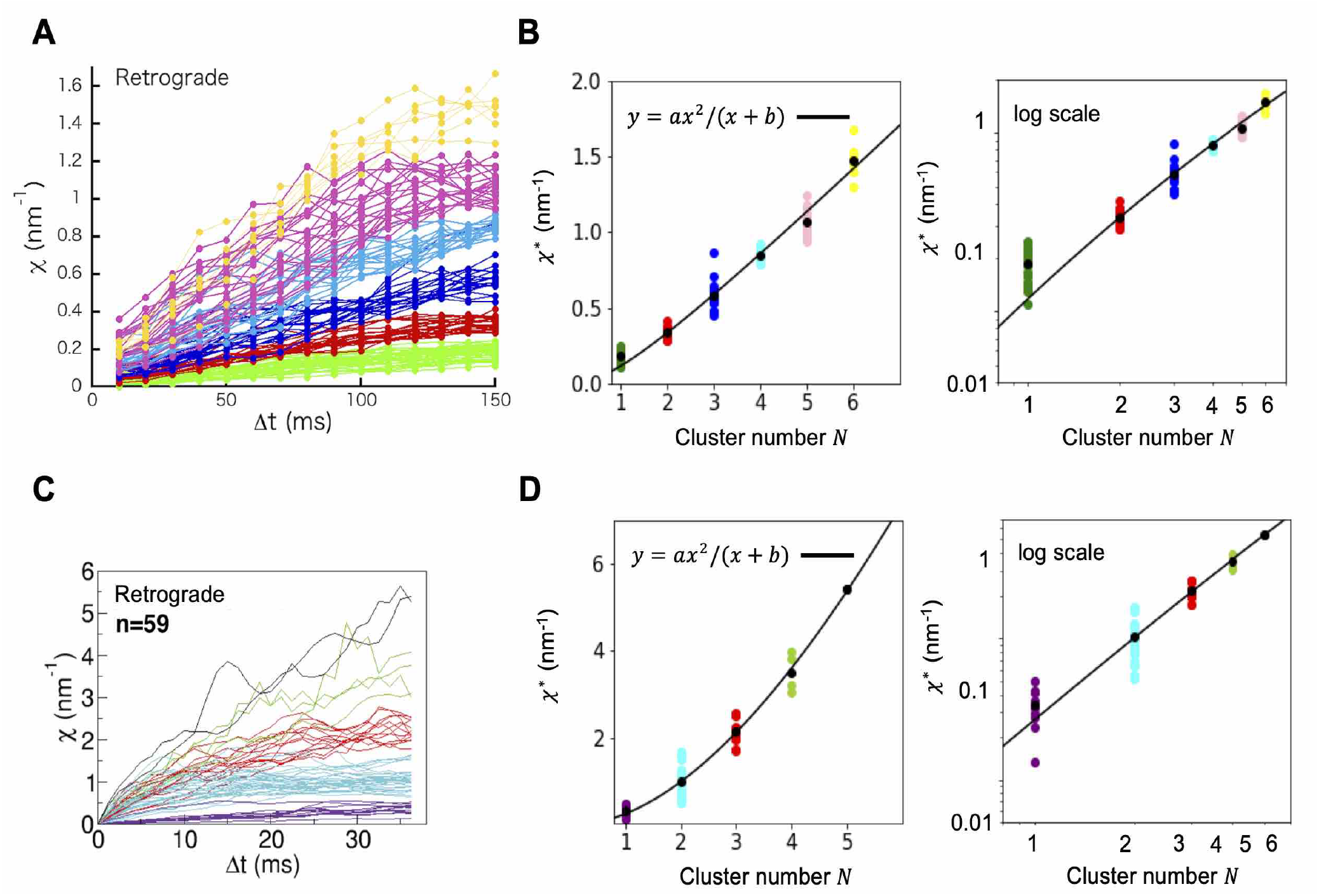
Measurement of χ for synaptic vesicle precursor transport by dynein. **A, B:** Measurement of χ for 116 cargo vesicles in mouse hippocampal neurons (Hayashi et al., 2021). We found six clusters. The convergent value, χ^∗^, of χ as a function of the cluster number *N*. The black line represents Eq. (1). **C, D:** Measurement of χ for 59 cargo vesicles (melanosomes) in zebrafish melanophores (Hasegawa et al., 2019). We found five clusters. The convergent value, χ^∗^, of χ as a function of the cluster number *N*. The black line represents Eq. (1).

## Conclusion

We introduce a novel physical measurement, termed χ, to provide insights into the mechanisms underlying KIF1A-associated neurological disorders, with particular emphasis on the role of the number of kinesin molecules. We suggest that the physical properties of axonal transport are governed not only by transport force and velocity, but also critically by the number of participating kinesin molecules. Prior research indeed suggests that variations in kinesin count can precipitate abnormalities in synapse formation (Chiba et al., 2019; Niwa et al., 2016). Hence, a comprehensive examination of the link between axonal transport aberrations related to the changes in the number of kinesin molecules associated to cargos and synaptic formation anomalies represents a crucial focus for future investigations. The potential implication of transport disorders extends to a spectrum of neurodegenerative diseases, including but not limited to Alzheimer’s disease, Parkinson’s disease, ALS, spastic paraplegia, Short Rib-Polydactyly Syndrome, and viral transport-related disorders. Our intention is to make a meaningful contribution to the clarification of molecular mechanisms underpinning neurological diseases associated with axonal transport, facilitated by our novel approach in physical measurements. Our novel approach, based on original physical measurement to estimate the number of kinesin molecules, can foster our understanding of the molecular perturbations in the axonal transport machinery and underpinning the development of neurological diseases.

## Acknowledgments

This research was conducted with the support of AMED under Grant Number JP18gm5810009, JST under Grant Number JPMJPR1877, the Nakatani Medical and Measurement Technology Promotion Foundation Development Research Grant, the KIF1A.org Mini Grant, and the Grant for Basic Science Research Projects of the Sumitomo Foundation.

